# A new statistic for efficient detection of repetitive sequences

**DOI:** 10.1101/420745

**Authors:** Sijie Chen, Fengzhu Sun, Michael S. Waterman, Xuegong Zhang

## Abstract

Detecting sequences containing repetitive regions is a basic bioinformatics task with many applications. Several methods have been developed for various types of repeat detection tasks. An efficient generic method for detecting all types of repetitive sequences is still desirable.

Inspired by the excellent properties and successful applications of the D_2_ family of statistics in comparative analyses of genomic sequences, we developed a new statistic 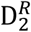 that can efficiently discriminate sequences with or without repetitive regions. Using the statistic, we developed an algorithm of linear complexity in both computation time and memory usage for detecting all types of repetitive sequences in multiple scenarios, including finding candidate CRISPR regions from bacterial genomic or metagenomics sequences. Simulation and real data experiments showed that the method works well on both assembled sequences and unassembled short reads.

## INTRODUCTION

The nucleotide sequences of most genomes contain many repetitive sequences of various types, such as short tandem repeats, interspersed repeats dispersed throughout the genome, or spaced repeats within a short region of the genome. Detection of repetitive regions in biological sequences is a basic problem in computational biology that has many applications. Repetitive sequences compose a large part of the genomes of eukaryotic organisms and have complicated features and functions (de Koning et al. 2011; Biscotti et al. 2015). They are also critical in prokaryotic genomes (Aras et al. 2003). Especially, a particular family of clustered spaced repeats in bacteria and archaea genomes called CRISPR or Clustered Regularly Interspaced Short Palindromic Repeats has been found to play important roles for the prokaryotes to protect themselves from viral attacks (Horvath and Barrangou 2010). This mechanism has been successfully applied by scientists to build the powerful technology of CRISPR/Cas9 that is now widely used to edit and engineer genes and genomes in any organism (Jinek et al. 2012; Cong et al. 2013; Mali et al. 2013; Hsu et al. 2014). Knowledge about repeats in genomes also helps to better solve computational tasks like assembly (Régnier and Chassignet 2016) and mapping (Misawa 2013).

The problem of detecting repetitive regions in genomes has been investigated for years, and many tools on repeat detection have been developed. For example, RepeatMasker (Smit et al.) and Censor (Jurka 2000) search the given sequence against repeat databases such as RepBase (Jurka 2000). These library-based methods are designed for specific species or specific repeat types. When the repeat type is not known in advance, or the species lack well-curated repeat database, de-novo repeat detection methods are better choices. Some *de novo* tools utilize a self-comparison strategy: TRF (Benson 1999) identifies repeats by scanning exact matches of small windows on genome; RECON (Bao and Eddy 2003) and PILER (Edgar and Myers 2005) extend local alignment algorithm to achieve repeat identification, while other *de novo* tools such as Tallymer detect repeats by counting the k-mer frequencies (Kurtz et al. 2008). Girgis et al. made a comprehensive review of genome-level repeat detection tools (Girgis 2015).

All genome sequences were obtained by assembling short sequencing reads. In many scenarios, it is beneficial if we can detect repeats from sequencing reads before they are assembled. For example, the existence of repetitive sequences is a major cause of difficulty for many assembly algorithms and a major aspect of the inaccuracy in reference genomes. For the types of short local repeats that can be observed within the length of a single read, it is desirable to rapidly screen reads with repeats so that reads with repeats and without repeats can be processed using different algorithms or parameter settings. For other scenarios such as finding clustered repeats like CRISPR that may have specific functions, the target patterns are usually very sparse in the genomes. Hence it is wasteful to assemble all sequencing reads into genomes or long scaffolds to find the repetitive regions. In such cases, being able to efficiently screen reads that contain repetitive regions will enable us to find the target repetitive patterns quickly without the heavy computation of sequence assembly.

Recent work on CRISPR provided a good example on the potential importance of studying repetitive sequence patterns in microbial genomes. CRISPR is a mechanism in many bacteria and archaea genomes for immune resistance against phages (Barrangou et al. 2007). The key characteristic of CRISPR is its repetitive structure in the microbial genome: a CRISPR cassette composed of a series of direct repeats of 21∼47bp each interspaced with spacer sequences of similar size that store fingerprint fragments of exogenous invading DNA sequences (Grissa et al. 2007). This mechanism has been adopted as a powerful technology for editing genomes and also as a tool to perturb the expression of genes (Dixit et al. 2016). Different microbial genomes may contain different CRISPR systems, which may have different properties for applications. Finding or designing new CRISPR systems that may have desirable advantages for some particular types of applications is of high scientific and technological value.

Microorganisms are the most abundant and diverse form of life on Earth. Natural selection has accumulated huge quantities of features in microbial genomes that are rich resources for discovering sequence elements that have the potential to be of engineering use, such as the CRISPR system. Most of the microbes have not been cultured or sequenced. Metagenomic sequencing provides an efficient way of sequencing the mixture of many microbial genomes (the microbiomes) of a microbial community. This provides a rich resource for mining useful genomic features. There have been several studies on discovering novel CRISPRs and other defense systems of microorganisms with the help of metagenomic data (Mangericao et al. 2016; Burstein et al. 2017; Doron et al. 2018).

Assembling the mixture of metagenomic sequencing reads into the component genomes or their long segments is a very challenging and computationally expensive task (Howe and Chain 2015). However, the typical length of sequencing reads is usually sufficient to include repeats in single reads if they are from CRISPR regions. As the majority of the metagenomic sequencing reads are not from CRISPR regions, it will be extremely efficient for finding potential CRISPRs from metagenomes if we can quickly pick up those reads with repetitive regions from the huge amount of unassembled sequencing reads.

Recently, there have been some new methods for assembling repeats or finding repeat motifs from short or long sequencing reads (Koch et al. 2014; Chu et al. 2016; Guo et al. 2018), filtering short reads containing tandem repeats (Misawa 2013) and assembling CRISPR candidates from metagenome sequences (Skennerton et al. 2013; Ben-Bassat and Chor 2015; Lei and Sun 2016). In this paper, we present a novel statistic 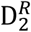 for the general purpose of detecting any types of repetitive regions efficiently in both computational time and memory, based on a variation of the D_2_ statistic for alignment-free sequence comparison. The statistic can be used as an efficient tool for both screening short reads with repetitive elements and identifying repetitive regions from long segments of genomes of full genome sequences.

The family of D_2_ statistics has shown impressive performance in revealing similarities between two sequences (Torney et al. 1990; Reinert et al. 2009). This inspired us to extend the idea of alignment-free sequence comparison to repeat detection in single sequences. In pairwise sequences comparison, word co-occurrences in two similar sequences lead to higher D_2_ scores, while non-similar sequences get lower scores. The basic idea of D_2_ statistic and its relatives is to measure sequence similarity by counting word matches between two sequences. Such a measure should also work for repeat detection in single sequences since there must be duplicated k-mers in a sequence with repeats. The count of self-matches of k-mers in a sequence can reflect the repetitiveness of the sequence, regardless of whether the repeat is tandem or interspaced.

Based on this idea, we developed a new variation of the D2 statistic for detecting the existence of repetitive regions within a sequence read or segment. We call the new statistic 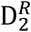 to mean “D_2_ for repeats”. We studied the statistical and computational properties of the new statistic. The computational complexity is linear, and the memory requirement is low. Experiments on simulated and real data showed that the statistic can distinguish sequences containing repetitive regions from those without repeats. The developed method based on 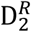 can be applied to many different scenarios for detecting repetitive sequences.

## RESULTS

### Experiments on simulated data

#### Results on simulation data

We generated 50,000 null sequences and 50,000 alternative sequences with the GC-rich model of p_a_ = p_t_ = 1/6, and p_g_ = p_c_ = 1/3. Each sequence is of 200-bp long. For the alternative sequences, we placed a repetitive region in each sequence at random positions. The repetitive region is composed of 4 replicates of a 12-bp repetitive unit (seed) interspaced with an 8-bp spacer between two consecutive seeds. This is done by using the seed sequence to replace the original sequence at the selected positions. To mimic more realistic situations, we introduced random errors to construct inexact repeats (30% of the repeat units have 2 letters being mutated).

We calculated the 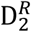 statistics with k-mer size from 3 to 15 on this dataset. Fig. 1 (a) and (b) show the histogram of 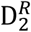 values for k=4 and k=7 among the positive and negative samples. In general, the value of 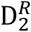 is larger in the positive samples than in the negative samples. For k=4, which is very small comparing to the length of the repetitive seed (12 in this case), the distributions of 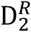 in the positive samples and negative samples have a big overlap. This is because that the occurrence of two or more instances of a 4-mer word in a random 200nt sequence segment is high, making the two classes not that separable based on 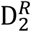 with 4-mers. For the case of k=7, although it is still smaller than the seed length, the two distributions become highly separable, with only a tiny overlap. The positive sequences can be distinguished from negative sequences with high accuracy if we choose a threshold at the intersection of the two distributions. The ROC curve of the method can be obtained by checking the true positive rates and false positive rates with different choices of the threshold.

**Figure 1.**
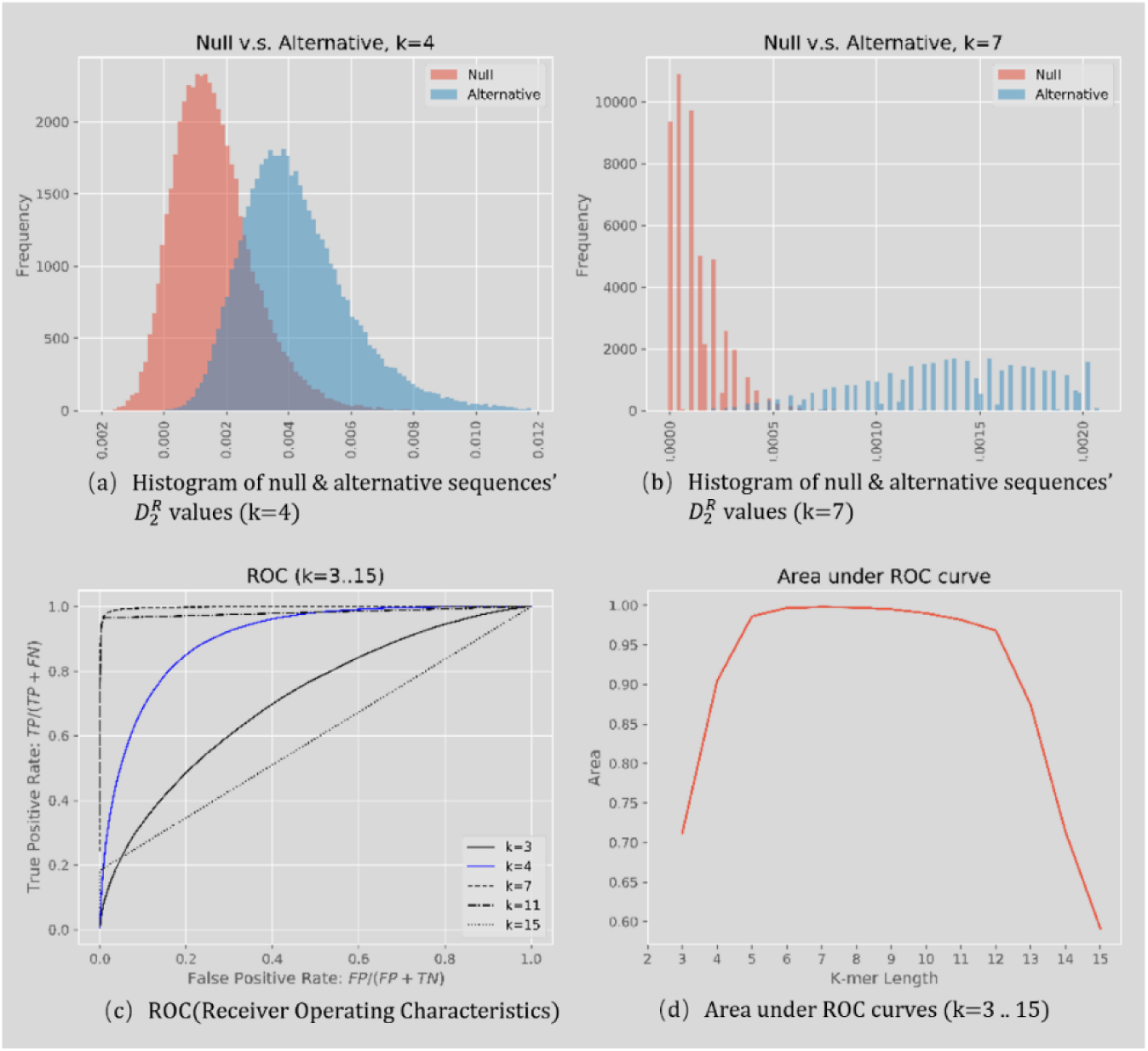
Results of repeat detection on simulated sequences: (a)(b) Histograms of 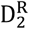 with k=4 and k=7. The null distribution (red) and alternative distribution (blue) get apart with proper k-mer size. The long tail of alternative distribution in (b) is truncated. (c)(d) The classification ROC curves and the area under the curves at different choices of k.

Fig. 1 (c) shows the ROC curves for k=3, 4, 7, 11 and 15. With k = 3, 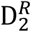 has some classification potential to distinguish positive from negative sequences but the classification accuracy is low, with AUC (Area Under the ROC curve) of only 0.71. The AUC increases to 0.84 at k=4 and reaches 0.99 at k=7. The performance starts to decrease when k=11, and classification accuracy drops steeply when k is larger than the length of the underlying repetitive element, and almost completely lost when k=15. Fig. 1 (d) shows the AUCs at different k-mer sizes from 3 to 15. Ideally, it may be expected that the method performs the best when the choice of k is equal to the length of the repetitive seed. But since errors were allowed in the repeats, the best performance is reached at k-mer sizes smaller than the seed length.

We performed extensive simulations with different parameter settings and different types of repeats, including spaced and tandem repeats with varied repeat unit length, both with noises and without noises. The results are given in the Supplemental Materials section 2. The observations are consistent with the above example.

These simulation studies show that the 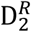 statistic is an efficient and powerful tool for identifying sequences with repetitive regions of different types. Curves are near the upper-left corner with both high sensitivity and high specificity at a wide range of choices of k, and the optimal performance can usually be reached when k is slightly smaller than the length of the underlying repetitive elements.

#### Comparison with RF

We also compared 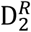 with the sequencing read screening tool RF (Misawa 2013) on the simulation data. We chose RF for comparison, rather than other well-known repeat finding tools such as RepeatMasker (Smit et al.) or TRF (Benson 1999), because it is a read-level repeat screening tool, while others detect repeats at a genome-level. RF suites the task of short sequence classification better. As RF can only detect tandem repeats, we only compare their classification performances in the tandem repeat scenarios.

Table 1 summarizes the results of two experimental groups of “strong” and “weak” signals. For the group of strong signals, sequences with long exact tandem repeats (12-bp exact repeat unit with 4 copies) were simulated. For the group of weak signals, sequences with short noisy tandem repeats were simulated (8-bp inexact repeat unit with 2 copies). In both group, 50,000 i.i.d sequences of length 200-bp were generated with the GC-rich model (p_a_ = p_t_ = 1/6, p_g_ = p_c_ = 1/3) to build negative samples. Equal number of positive samples were built by tandemly inserting repeat units into the random sequences. We chose multiple thresholds for RF and 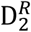 and compared the classifiers’ accuracy, precision, recall, and time consumption. We used 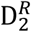 (k-mer size=7) for the strong signal group and 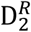 (k-mer size=8) for the weak signal group because they have the highest AUC among all k-mer sizes.

**Table 1.** Comparison of the performance of 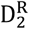 and RF

| Experiment | Method | Parameters | Accuracy | Precision | Recall | Time |
| --- | --- | --- | --- | --- | --- | --- |
| Strong signal | RF | Threshold=20 | 1.000 | 1.000 | 1.000 | 9,075 ms |
| Strong signal | $D_2^R$ | Threshold=2.2e-3, K=7 | 1.000 | 1.000 | 1.000 | 1,622 ms |
| Weak signal | RF | Threshold=7 | 0.563 | 0.694 | 0.224 | 11,117 ms |
| Weak signal | RF | Threshold=10 | 0.563 | 0.694 | 0.224 | 10,029 ms |
| Weak signal | RF | Threshold=11 | 0.533 | 0.726 | 0.106 | 10,996 ms |
| Weak signal | RF | Threshold=13 | 0.507 | 0.757 | 0.022 | 11,304 ms |
| Weak signal | RF | Threshold=15 | 0.502 | 0.812 | 0.004 | 12,006 ms |
| Weak signal | $D_2^R$ | Threshold=3.0e-6, K=8 | 0.755 | 0.700 | 0.893 | 843ms |
| Weak signal | $D_2^R$ | Threshold=3.6e-5, K=8 | 0.755 | 0.700 | 0.893 | 1,863ms |
| Weak signal | $D_2^R$ | Threshold=1.0e-4, K=8 | 0.706 | 0.774 | 0.582 | 1,634 ms |
| Weak signal | $D_2^R$ | Threshold=1.5e-4, K=8 | 0.629 | 0.819 | 0.330 | 1,455 ms |
| Weak signal | $D_2^R$ | Threshold=3.0e-4, K=8 | 0.519 | 0.864 | 0.044 | 1,636 ms |
| Weak signal | $D_2^R$ | Threshold=5.0e-4, K=8 | 0.502 | 0.836 | 0.004 | 1,535 ms |

Two experiment groups were used, one with strong signal and one with weak signal. It is more difficult to distinguish repetitive reads in the weak signal group. Both $D_2^R$ and RF perform well in the strong signal group. In the weak signal group, $D_2^R$ outperforms RF. And $D_2^R$ is much faster in all cases.

Classification of sequences in the strong signal group is an easy task for the two methods. They can both reach perfect performance (Accuracy = 100%) at proper parameter settings. For the weak signal cases, 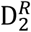 performs better in terms of the receiver operating characteristic (Fig. 2) and has higher accuracy in general. Both RF and 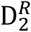 were implanted in Java and our program is about 9 times faster than RF.

**Figure 2.**
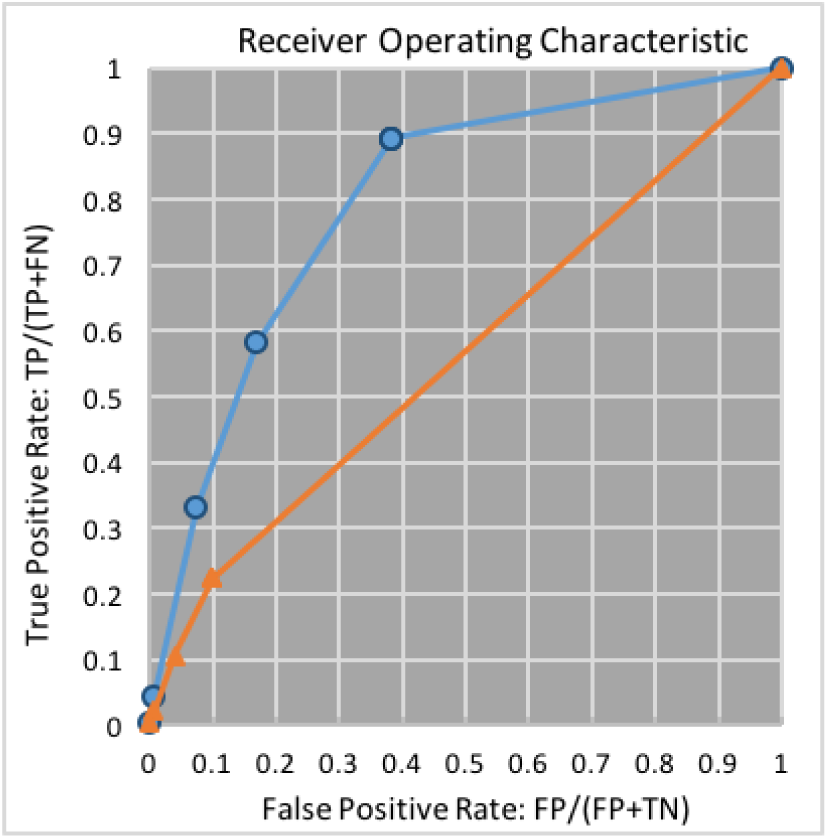
Comparison of the ROC curves of 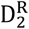 (in blue) and RF (in orange) in the weak signal experiments. The points of 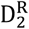 experiments are closer to the top-left corner compared with RF, which means 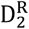 has higher sensitivity and specificity.

### Experiments on real data and automatic choosing of the threshold

#### Results on a prokaryotic genome

We conducted genome-level repeat detection experiments on a prokaryotic genome as an example. The genome we used was *Archaeoglobus fulgidus DSM 4304*, downloaded from https://www.ncbi.nlm.nih.gov/nuccore/NC_000917. The genome length is 2,178,400 bps. The task of the experiment was to find candidate CRISPRs in this genome to illustrate how 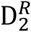 works on long sequence scaffolds or full genomes, and how the threshold can be decided automatically in an unsupervised manner.

Firstly, we counted the nucleotide frequencies on this genome, which are p_a_ = 0.2580, p_t_ = 0.2562, p_g_ = 0.2438, p_c_ = 0.2420. Then we set the sliding window size=1,000 bps and used Algorithm 2 to scan the genome with k=7. Fig. 3 (a) shows the scanned 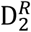 values along the genome. Three major peaks in the plot are identified, indicating three regions that contain strong local repetitive patterns in the genome. They are the candidate CRISPR regions. Fig. 3 (b) gives the zoom-in of the three peaks. Each peak has a plateau of high 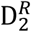 values. We find that these three peak regions correspond to the three confirmed CRISPR structures that had been recorded in CRISPRdb[19], at locations (148, 4,213), (398,369, 401,590), and (1,690,930, 1,694,181), respectively. We can also see that the plateau part of each peak shows some oscillation in the 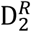 value, which is in accordance with the CRISPR’s interspaced repeat structure.

**Figure 3.**
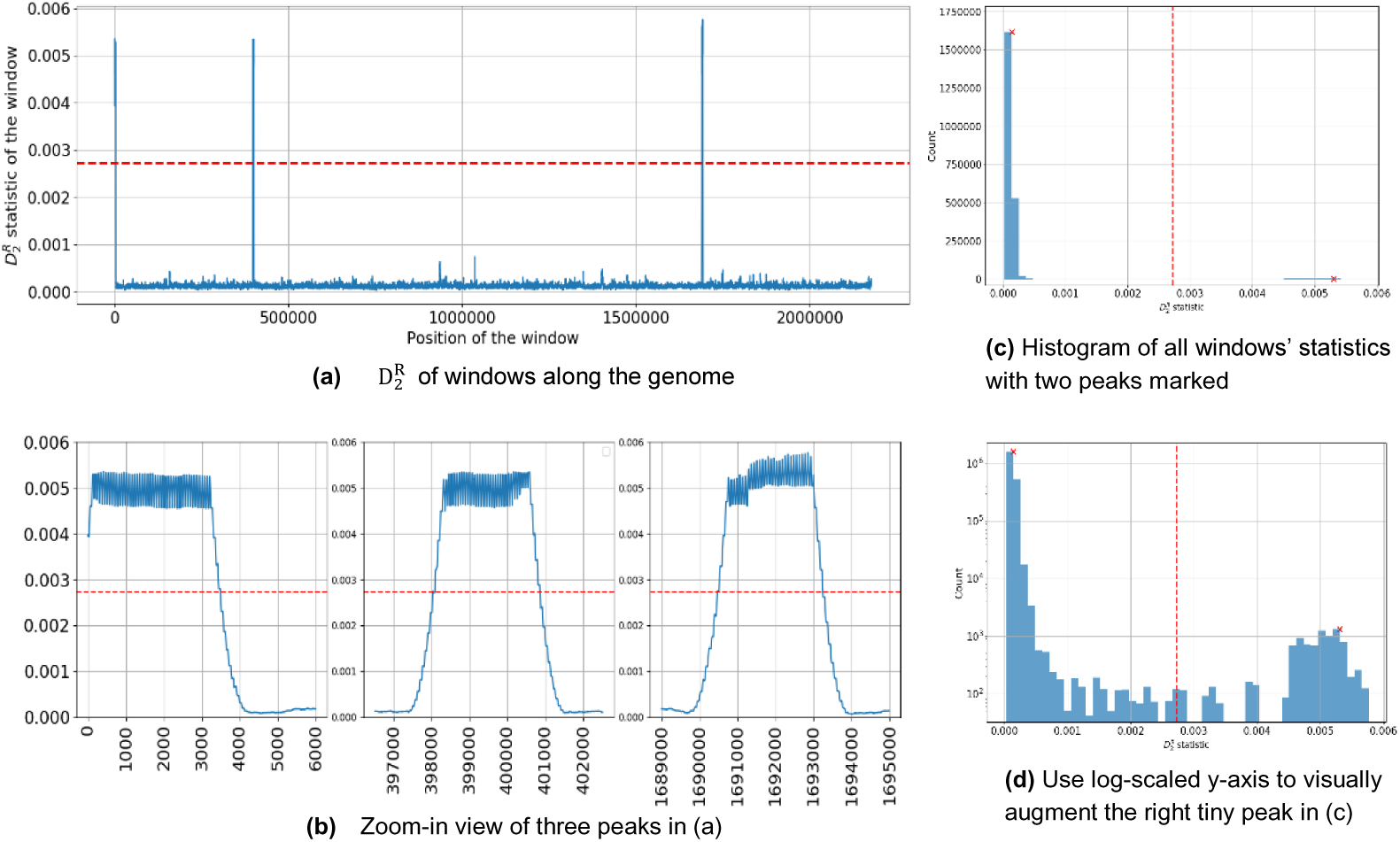
Results of repeat detection on real genome. (a) 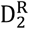 values of 1kb-windows along the genome. (b) Zoom-in of the three peak regions. The red dashed line shows the threshold value determined by the unsupervised method. (c) The histogram of the 1kb-windows’ 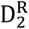, with the position of the threshold value (0.002723) marked with red dashed line. The non-repetitive windows contribute to the large peak around 0, while a small number of repetitive windows contribute to the tiny peak around 0.005. (b) The log-scale plot of the histogram which can show the tiny peak at high 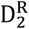 value.

**Figure 4.**
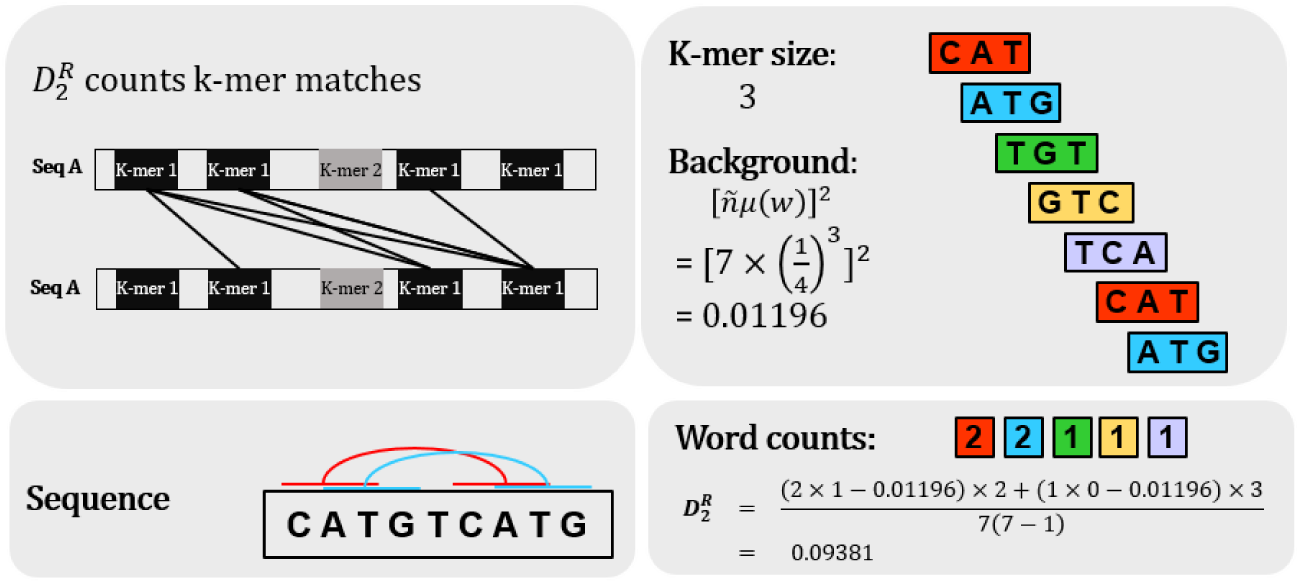
Matches within a single sequence. 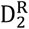 counts k-mer matches within a single sequence. The definition of 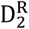 amplifies the signal of duplicated k-mers and restrains the signal of k-mers appearing only once.

**Figure 5.**
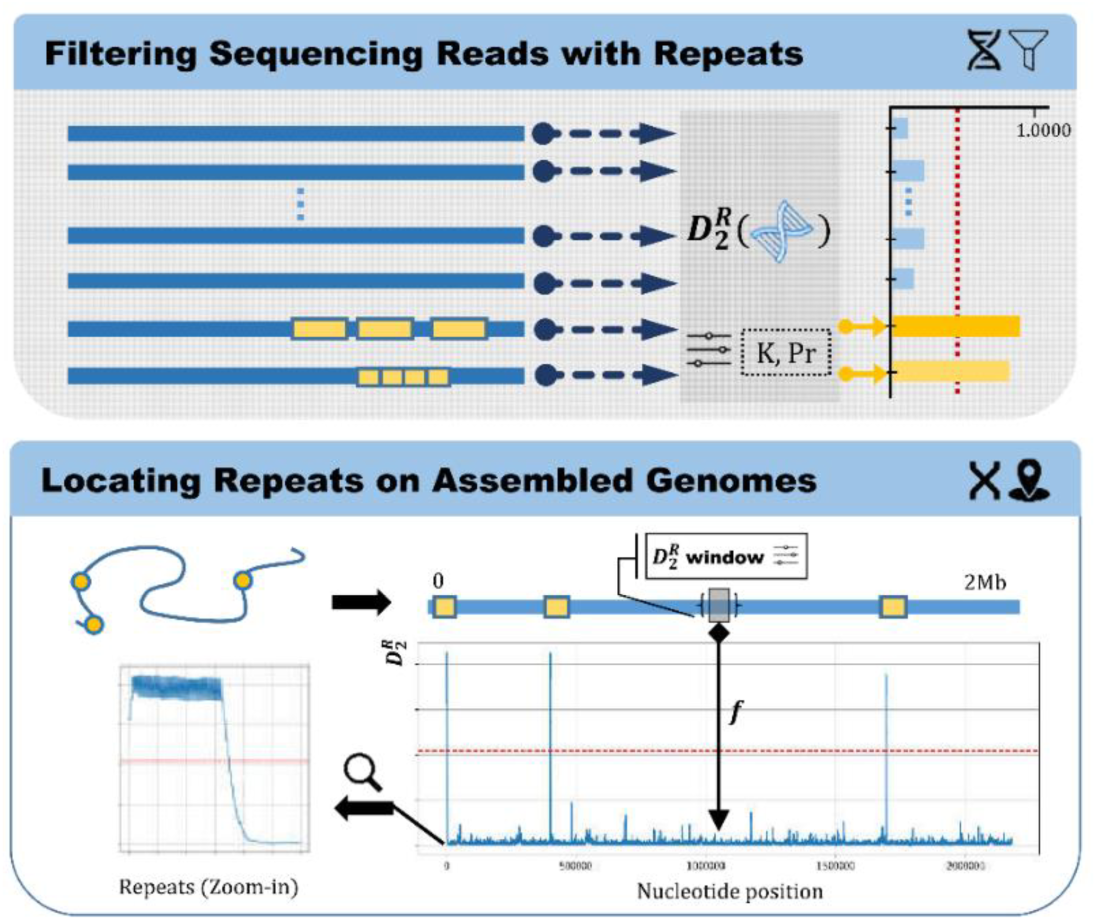
Two scenarios of repeat detection with 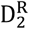: 1. The 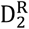 statistic evaluates the repetitiveness of each read. With proper parameters and threshold values, reads containing full or part of repetitive structure (tandem repeats or interspaced repeats) can be detected. 2. The 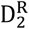 statistic locates the repetitive structure by scanning a long sequence segment by sliding windows. An implemented window-updating algorithm can scan a typical bacteria genome within less than a second on a personal computer. In these two scenarios, 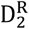 plays a role as a filter with configurable parameters (k-mer size and background sequence probability model, shown as “K” and “Pr” in the figure), which filters out the non-repetitive sequences and keep the repetitive ones.

**Figure 6.**
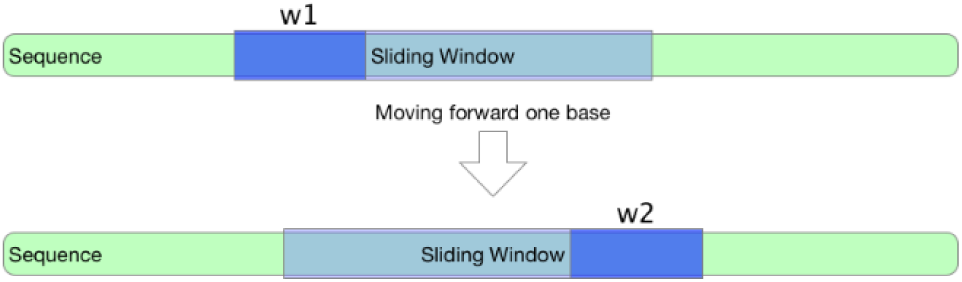
A sliding window moving one letter forward on the genome. In each movement, one new word is appended to the tail of the sliding window and one word in the front is discarded.

The distribution of 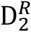 values of the sliding windows on the genome suggests a way to automatically decide on the threshold. As we can see, the 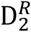 values at the CRISPR regions are much higher than the other values along the genome. When we draw the histogram of all 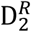 values as in Fig. 3(c), most values are distributed at the small-value end of the histogram, but there is a tiny peak at very large values. This bimodal nature of the distribution can be seen more clearly on the log-scaled histogram shown in Fig. 3(d). The threshold can be set between the two peaks at 0.002723 as shown in Fig. 3(d).

When we use this method to scan a new genome, if distributions of this style are observed, we can infer that there must be some regions with strong repetitive patterns, and we can also easily choose a mid-point between the two peaks as the threshold for the detection. Furthermore, when we apply 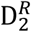 on unassembled sequencing reads in genomic or metagenomic sequencing data, we can also use a similar strategy to infer the existence of repetitive sequences and to help choosing the threshold.

As a practical concern in real applications, since usually the majority of sliding window locations along the genome and the majority of the unassembled sequencing reads do not contain the target repetitive regions we are looking for, it can be expected that the tiny peak at high 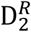 values can be very weak and are easily missed. Zoom-in view of the high-value end of the histogram or using log-scale can be helpful if the histogram is to be manually studied, as shown in Fig. 3(d). To make the analysis more automatic, we employed a peak-detection algorithm to find peaks in the histogram. It is based on the continuous wavelet transform, provided by the python *scipy* package originally developed for mass spectrometry data (Du et al. 2006). Applying this method on this histogram of Fig. 3(c), two peaks were identified as marked by the red crosses on the plot, which are also shown in Fig. 3(d). The two peaks are at 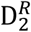 value positions of 0.000141 and 0.005306, respectively. We chose the mid-point position 0.002723 between two peaks as the threshold. The red dashed lines drawn in Fig. 3 show the location of the threshold on the 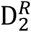 value axis. The three CRISPR regions were captured accurately with using this cut-off.

#### Comparison with other methods

We also compared our method with two other repeat finding tools on this genome scanning task (Table 2). We considered a CRISPR detection tool CRT (Bland et al. 2007) as well as a recently-published genome-level repeat detection tool Red (Girgis 2015). All three methods successfully detect the CRISPR region but our program is the fastest in speed. Parallel optimizations like multithreading have not been implemented in the current version of our program. Therefore, it has a large space to be further accelerated through parallel implementation.

**Table 2.** Efficiency comparison of the genome scanning task

| Method | Parameters | Average Time |
| --- | --- | --- |
| $D_2^R$ | Window size = 1000,<br>k-mer size = 7 | 563 ms |
| CRT<br>(2007) | MinNumberRepeats=3, MinRepeatLength=15,<br>MaxRepeatLength=48, SearhWindow=8,<br>MinSpacerLength=15, MaxSpacerLength=48 | 812 ms |
| Red<br>(2016) | Default parameters | 6,127 ms |

CRT is specially designed for CRISPR region detection in genomes. It accepts assembled genomes and outputs identified CRISPR regions with their locations and the putative repeat/spacer units. Red detects repeats of all types from the input genome and annotates the location of repeats in output file. Our algorithm scans the given genome and returns a file containing $D_2^R$ of all sliding windows. The running time shows that $D_2^R$ is efficient in detection of repeats of various types. The genome used in this experiment is *Archaeoglobus fulgidus DSM 4304*. This comparison is conducted on a PC with Intel core i7 6700 CPU, 8GB memory, and on Windows 10 operating system.

## DISCUSSION

We proposed a new statistic 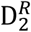 that can be used to detect the existence of repetitive regions in a short or long sequence with high accuracy. We developed algorithms for calculating the statistic and using it for repeat detection with high memory-and computation-efficiency. It is a very handy tool for many applications that involve the scanning or detection of repetitive sequences, such as finding CRISPR-like repetitive regions from bacteria genomes or from the massive unassembled sequencing reads in metagenome data.

Experiments showed that the statistic has good performance on different scenarios and for different types of repetitive patterns. But the theoretical characterization of its mathematical properties is not fully available yet. The choice of the optimal k of the k-mer size depends on several factors such as the total length of the sequence, the length of the repetitive unit and the level of noise in the repeats. The theoretical choice of the optimal threshold of 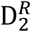 to discriminate sequences with and without repetitive regions is still not available. Experiments show that there is a large range of suboptimal choices of k that can result in almost the same performances. And in practical applications, as shown in our simulation and real data experiments, the distinction of 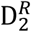 values of sequences with and without repeats is large and the threshold can be easily determined through known data or be decided automatically in an unsupervised manner through the distribution of 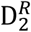 values on the data. We also developed a method to automatically determine the threshold in this manner.

## METHODS

### From similarity to repetitiveness

The D_2_ statistic counts the number of k-mer matches between two sequences. Given two sequences A = A_1_A_2_…A_n_ and B = B_1_B_2_…B_m_ over the alphabet 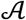, the original D_2_ statistic is defined as (Torney et al. 1990; Lippert et al. 2002):
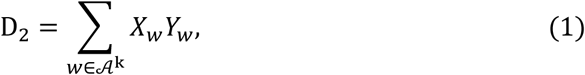
where 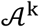 is the set of all k-mers or k-tuple words (words of k letters from the alphabet 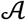), and X_w_ and Y_w_ are the numbers of occurrences of the k-tuple word w in sequences A and B, respectively. For pairwise sequence comparison, this product of k-mer counts can be viewed as the number of k-mer matches between the two sequences. More matches indicate higher similarities between the two sequences.

We extend this idea to the single sequence case: using self-similarities of a sequence to measure its repetitiveness. Consider a k-mer word w appearing X_w_ times in a sequence S of length n. The number of self-matches within S is 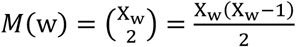.

The total number of self-matches of all k-mers in the sequence is a rough reflection of the repetitiveness of the sequence. It is obvious that the longer a sequence is, the higher the chance is that there are more self-matches. We need to normalize this count by the maximum possible number of matches given the sequence length. We define 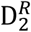 as the normalized sum of counts of all k-mer matches:
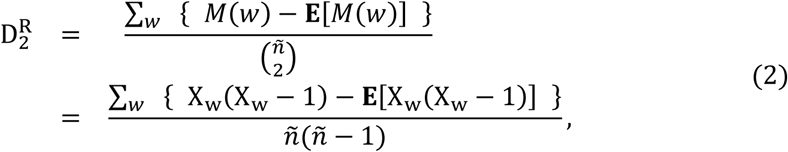
where *ñ* =*n* − *k* + 1 represents the number of all k-mers in the sequence S. In the numerator, we subtract the expectation of random self-matches by chance as background from the observed sum of self-matches. In equation (6), we will show that **E**[X_w_(X_w_ − 1)] can be approximated by [*ñ*μ(*W*)]^2^ where μ(*W*) is the expected word count of w under the assumption that X_w_ is Poisson distributed. For a sequence of length n, the maximal number of matches that the *ñ* k-mers may form is 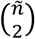. So we employ 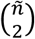 as the normalization term to measure a relative level of match count.

With this definition, we can infer that the values of 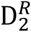 will be high for sequences that contain repetitive regions no matter if the repeats are tandem or interspaced if the k is properly chosen, since the k-mers within the repetitive elements will occur more times than their expected number of occurrence by chance. For sequences without repeats, the number of a k-mer’s occurrence will be around the expected number so that 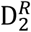 will be small. Hence, we can distinguish two types of sequences by setting a threshold on the 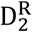 value.

Besides the presented form of 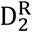, there can be other ways to define the statistic to implement the idea. For instance, Euclidean distance is often considered when evaluating the distance between two k-mer content vectors. It is possible that the norm of the k-mer content vector can also reflect the repetitiveness of a sequence. We have experimented with some other alternative definitions of the statistic. Results showed that those forms can also be used to distinguish sequences with and without repetitive regions, but their performance were not as good as the one defined by (2). The results are presented and discussed in Supplemental Materials section 4.1 and 4.2.

### 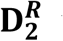 under the i.i.d. background model

To calculate 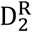, we need to first calculate or estimate **E**[X_w_(X_w_ − 1)] for the background model of sequences without repeats. It depends on several factors including sequence length, k-mer size, and the probability model for the background sequences. For simplicity, we assume each letter in the DNA sequence (and therefore in each k-mer word) is independently and identically drawn from the alphabet {a, g, c, t} with a certain probability (p_a_, p_g_, p_c_, p_t_). Although real genomes can have more complicated underlying sequence structure, this model can capture the basic nucleotide composition information and performs well in our experiments.

Considering a word w = (w_1_,…, w_k_) of length k, let II_i_(w) = {1, 0} be the indicator for w occurring at position i in the sequence. The word occurrence probability μ(w) is the product of letter occurrence probability since we assume each letter is independently drawn from the alphabet:
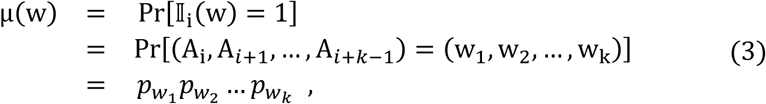
II_i_(w) is a Bernoulli random variable and its sum 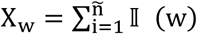 over all possible ñ = n − k + 1 positions in the sequence counts the word occurrence in the sequence. A closed form of the mean and variance of X_w_ can be found in (Waterman 1995):
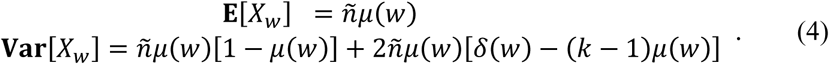
The δ(w) is an overlapping coefficient
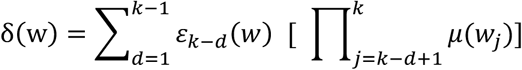
that measures the word’s ability to overlap itself, where ε_u_ (*W*) is an overlapping indicator that equals to 1 if the *u*-letter prefix is the same as the *u*-letter suffix in the word *w*, such as “gt” is both the prefix and the suffix of the word “gtacgt”, for example, and otherwise 0.

As a corollary, the mean of *X*_*W*_(*X*_*W*_ − 1) is:

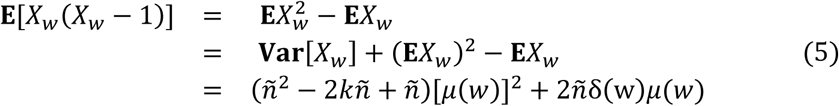

The derivation of variance of *X*_*W*_(*X*_*W*_ − 1) is non-trivial and a simple closed form has not been available. Some algorithms have been proposed to compute the probability generating function of *X*_*W*_, which helps to derive the moments of *X*_*W*_ and the variance of *X*_*W*_(*X*_*W*_ − 1), but these methods are word-specific and cannot yield a closed form variance (Ribeca and Raineri 2008; Nuel 2010).

In practice, we adopt an approximation of **E**[*X*_*W*_(*X*_*W*_ − 1)] by assuming the word count statistic follows a Poisson distribution, i.e. *X*_*W*_∼Poisson(*λ* = *ñμ*(*w*)). This assumption is feasible because as *ñ* increases and k is relatively large, the asymptotic distribution for *X*_*W*_ is a compound-Poisson distribution. With this assumption, the mean of *X*_*W*_(*X*_*W*_ − 1) becomes:
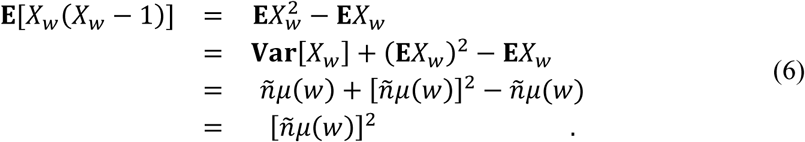
We can see that the expectation of *X*_*W*_(*X*_*W*_ − 1) calculated this way keeps the dominant term related to n^2^ in equation (5), and ignores the terms related to n^2^. With this simplification, we adopted the formula for 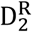 statistic for practical use as:
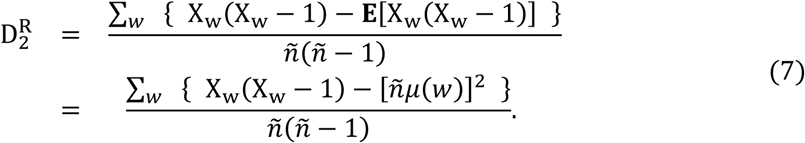
Experiments also show that this definition using the simplification of (6) performs equally well on the classification performance with that using the more time-consuming calculation of (5). (see experiments in the Supplemental Materials section 4.2)

Similar to the definition of 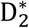 for the comparison of two sequences (Reinert et al. 2009; Wan et al. 2010), we also explored an alternative statistic that divides each term of 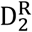 by the standard deviation of X_w_(X_w_ − 1). However, though this alternative statistic adds power in some experiments when the k-mer is short, it does not significantly improve the performance when k-mer is longer and even fails in some situations. (Supplemental Materials section 4.2) Therefore, we only present the results using 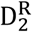 in the main text while the results using other alternative statistics are presented in Supplemental Materials.

### The repeat detection algorithm and its complexity analysis

We developed algorithms to use 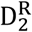 to detect repetitive regions for two typical scenarios. One is the screening of unassembled sequencing reads or short sequence segments that contain repetitive regions. The other is detecting and locating regions with local repeats in a genome or very long segments of assembled sequences.

Once the 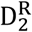 of a sequencing read or of a sliding window is calculated, it is straightforward to make a judgment on whether the current read or window contains repetitive regions by setting a proper threshold between 0 and 1. We present three threshold selection methods in the latter section. In some scenarios, however, it is more efficient to rank the candidates by the 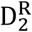 values and select the top candidates for additional studies instead of setting a fixed threshold (e.g. annotating repeats of unknown types in a new genome).

#### Computing 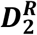 in a short sequence segment

The algorithm we developed for computing 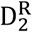 in a short sequencing read or segment is:

|  |
| --- |
| **Algorithm 1:** Compute $D_2^R$ in a single sequence |
| **Input:** A sequence $S$ of length $n$,<br>K-mer size $k$,<br>Base frequency vector* $(p_T, p_C, p_G, p_A)$ |
| **Output:** $D_2^R$ |
| **Step 0:** Initialize a k-mer occurrence hash table** $HashCount[ ]$:<br>The key $w$ is a k-mer and its value $X_w$ is the raw count of its occurrences;<br>Initialize $D_2^R \leftarrow 0$ |
| **Step 1:** Iterate all starting positions of k-mers on $S$ from 1 to $n-k+1$ to count k-mer occurrences.<br>**for** $i = 1, 2, \dots, n-k+1$ **do**<br>$HashCount[ \text{the k-mer at position } i ] ++$;<br>**end for** |
| **Step 2:** For each key-value pair in $HashCount[ ]$, convert the raw count to the mean-adjusted match count, as the numerator of (7).<br>**for** $w, X_w$ in $HashCount[ ]$ **do**<br>$RawCount \leftarrow X_w(X_w - 1)$<br>$\mu(w) \leftarrow p_{w_1} p_{w_2} \dots p_{w_k}$<br>$Mean \leftarrow (n - k + 1)^2 \mu(w)^2$<br>$MeanAdjustedCount \leftarrow RawCount - Mean$<br>$D_2^R \leftarrow D_2^R + MeanAdjustedCount$<br>**end for** |
| **Step 3:** Normalize $D_2^R$ as defined in (7),<br>$D_2^R \leftarrow D_2^R / [(n - k + 1)(n - k)]$<br>**return** $D_2^R$ |

\* Base frequency vector can be set according to knowledge or estimated from sample sequences.
\*\* The hash table is implemented with C++11 Standard Template Library's *unordered\_map*.

To compute the value of a single sequence of length *n*, in **Step** 1, the algorithm first lists all (*n-k*+1) k-mers in the sequence and count their occurrence with a hash table. This process takes O(*n*) time as inserting or updating a hash table take a constant time. After the k-mer counting step, we need to sum all O(*n-k*+1) = O(*n-k*) mean-adjusted match counts in **Step 2**, each costing O(*k*) time. The total time consumption is O(*k*(*n-k*)) = O(*n*) because the *k* is a small constant. The space consumption is O(*n-k*+1) = O(*n-k*) = O(*n*). Both time and space complexity are very low because lengths of sequencing reads are usually short.

As the computation of 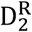 for each sequencing read is very light and is independent of other reads, we can compute the 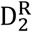 of massive sequencing reads in parallel with high efficiency when searching for reads with repetitive regions from massive amount of raw sequencing reads, such as in the scenario of looking for candidate new CRISPR elements from metagenome sequencing data.

#### Computing 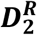 for sliding windows on a genome

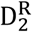 can also be used to scan for repetitive regions in a whole genome or a long sequence segment. Consider a sliding window of length *G* moving from one end of the genome sequence to the other end, the 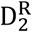 value of the sliding window at each position reflects the repetitiveness of each local region in the genome.

Repeatedly calculating 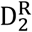 of each sliding window on the genome as an independent short segment wastes a lot of time, since the sliding windows overlap. Therefore, we designed an algorithm to update the 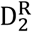 value when sliding window moves one base forward. The computational cost of the updating is of constant time.

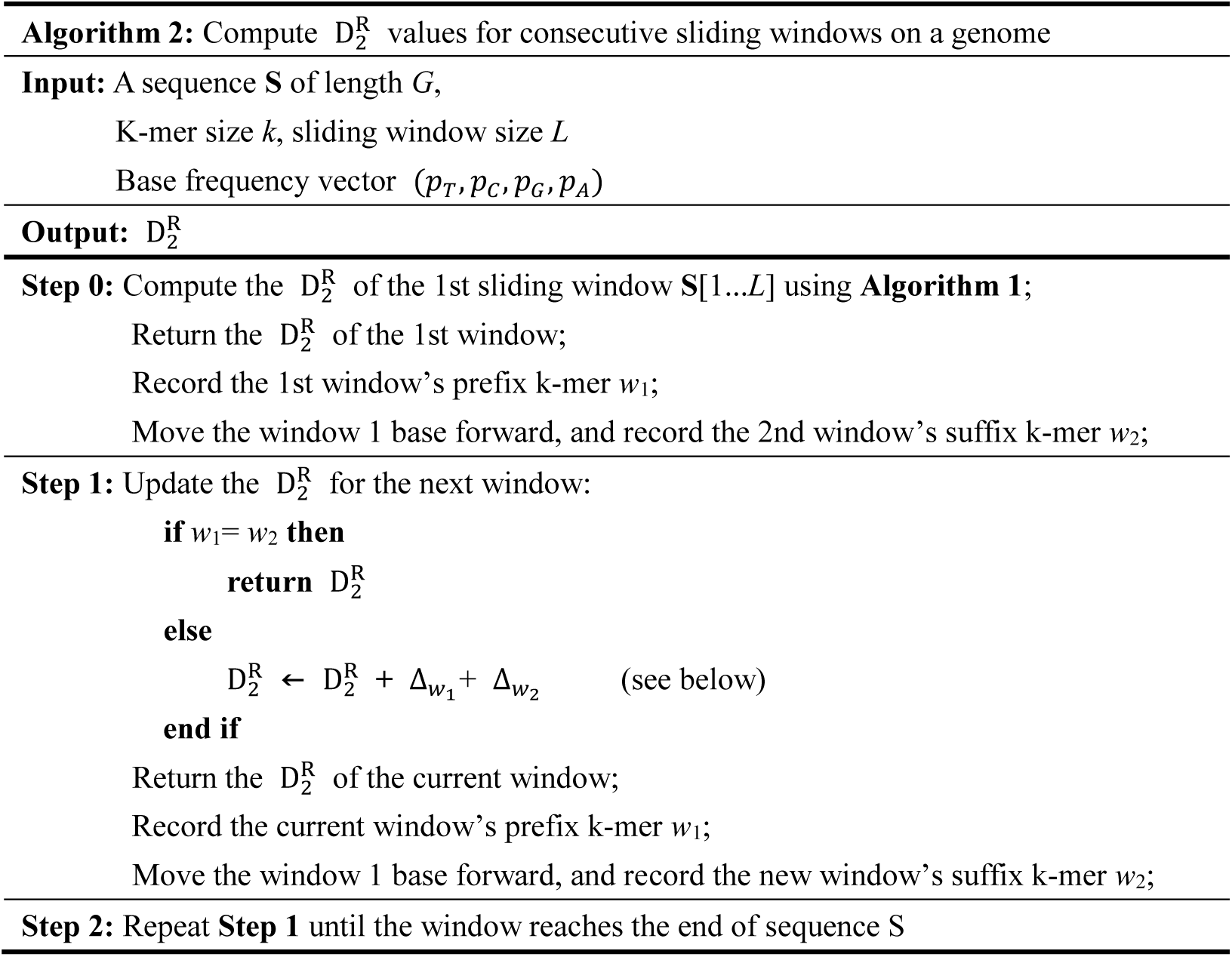

The details of updating 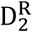 in **Step 1** are as follows. Let w_1_ be the first k-mer (the prefix) of the sliding window before movement, and w_2_ be the last k-mer (the suffix) of the sliding window after movement. Moving forward 1 base does not affect most of the k-mers’ frequency in the genome except for w_1_ and w_2_.

If w_1_ = w_2_, the statistic remains the same after movement because the k-mer content of the two sliding windows does not change.

If w_1_ ≠ w_2_, the new counts for the two words are 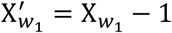, and 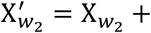 1, respectively. Let 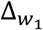 and 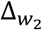 be the change of 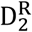 w.r.t w_1_ and w_2_ after the movement, we have
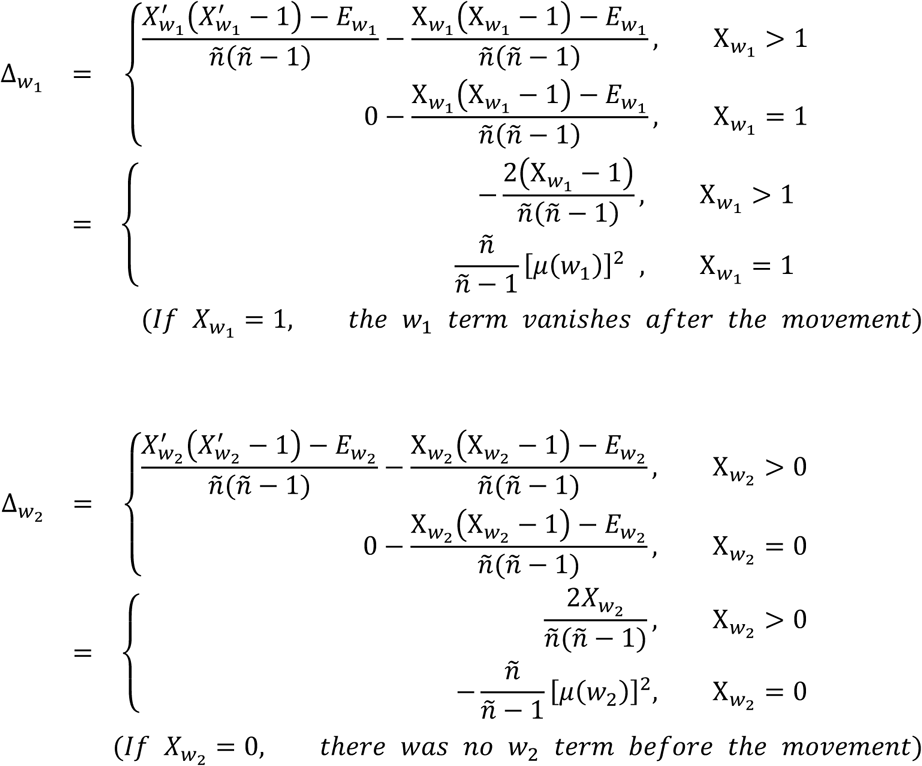
where *E*_*W*_ = **E**[*X*_*W*_(*X*_*W*_ − 1)] = [*ñμ*(*w*)]^2^. Given the above rules, we can update the value of 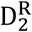 with 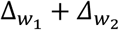 in O(*k*) time.

Suppose we have a genome of length *G*, the total time consumption for scanning the whole genome is O(*kG*) = O(*G*). This genome scanning process can be accelerated by starting multiple scanning head on the genome in parallel.

In summary, the repetitive region detection algorithm in both scenarios has a linear time complexity (proportional to sequence size) and a linear space complexity (proportional to read/window size).

### How to set parameters for the algorithm?

#### K-mer size

Choosing proper k-mer size assists the algorithm to achieve the state-of-the-art performance. One advantage of 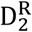 is it is not very sensitive to the selection of k. Shown in the simulation results in Fig. 1 (d) and Supplemental Fig. 2∼6, a wide range of k works nearly equally well for these cases.

When we have expectations about the type of repeats we are looking for, choosing a proper k with lowest classification error or other criteria based on the simulation result is not difficult. Typically, a classifier with k-mer sizes slightly smaller than the length of the repeat unit is robust to the noise occurred in inexact repeats.

#### Background sequence model

For simplicity, we used an i.i.d. model to evaluate the k-mer occurrence probability in this article. The occurrence probability of each base (a, g, c, t) can be estimated by counting the frequency in the real sequences.

However, the background sequence model may not be the best solution to represent the background non-repetitive sequence. If we hope to ignore the repetitive patterns caused by the sequence model and only report the real significant sequence, an i.i.d. model might over-simplified the problem. One way to alleviate this is to use a Markov chain to model the sequences and using BIC to determine the order of the Markov chain.

### How to choose threshold for 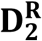 ?

Knowing the theoretical distributions of 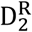 under the null and alternative model would guide the threshold selection because we would be able to compute the threshold with minimum classification error. However, a closed-form of the theoretical distributions are not available yet, and the threshold for detecting repetitive sequences has to be set heuristically. We suggest three ways for choosing the threshold – a supervised method, an unsupervised method, and a Q-value method.

#### Supervised method

One way is to decide the threshold by choosing some training sequence samples with and without repetitive regions and select the threshold that separates the distributions of 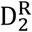 of the two classes with minimum classification error. For example, for the task of searching for potential new CRISPR sequences from metagenomic sequencing reads, one can use a set of known CRISPR sequences to compose the positive training set. The negative training set can be either composed by sequencing reads that are manually checked to have no CRISPR-like repetitive patterns, or composed by a random collection of reads from all the metagenomics sequencing reads. Given the fact that CRISPRs compose only a very small proportion of a metagenomics data set, the random collection of reads can also guarantee that most, if not all, of the samples are negative. This can save the effort of manually selecting negative training data.

#### Unsupervised method

A more automatic way for choosing the threshold is to decide it in an unsupervised manner. The idea is, if we can assume that the target sequences in a dataset are composed of sequences with repeats and sequences without repeats, then with an appropriate k, the distribution of 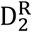 of all sequences in the dataset will be of dual modal. One can use the trough position as the threshold. This idea will be illustrated in our experiments on real genome data in the next section.

#### Q-value method

If the distribution is not of clear dual modal pattern, for example when there are multiple types of repeats in the genome, it is still feasible to determine the threshold by controlling the FDR (False Discovery Rate) using simulation data. One can first simulate a large number (e.g. 100,000) of sequences under the null model of length L, and calculate their 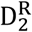 values. For each candidate sequence’s 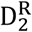 value from the real data, we can estimate its p-value by the fraction of null sequences with 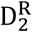 greater than the one from real data. In this way, we can get the empirical distribution of p-values for all the candidate sequences, and then will be able to get the Q-value (Storey 2003) for each candidate sequence from the real data. The threshold for 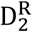 can be determined by setting it to control the FDR level.

### Sequence Simulation

We conducted a series of experiments on simulation data to study the performance of the proposed method. We generated null sequences by drawing letters from a given i.i.d. nucleotide distribution model, and generated alternative sequences by inserting multiple copies of seed substrings into null sequences. We take the null sequences as “repeat-free” and the alternative sequences as sequences with repetitive regions. There could be some sequences under the null model that may contain some repetitive elements by chance, but the probability is low for long repetitive elements. We took all repeat-free sequences as negative samples and the alternative sequences as positive samples, and studied the performance of the 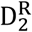 method to distinguish the two classes under different settings of the simulation and of the parameters in the method. Details of simulation procedures are described in Supplemental Material section 1.

## DATA ACCESS

The codes of the proposed method are available for free academic use at https://github.com/chansigit/D2R_codes

## ACKNOWLEDGEMENT

We thank Prof. Gesine Reinert of the University of Oxford for her helpful discussions. This work was supported by NSFC grants 61721003 and 61673231. F.S. and M.S.W. were supported by the 111 Project (Grant No. B18015), Shanghai Municipal Science and Technology Major Project, and US National Science Foundation (DMS-1518001).

## Conflict of Interest

none declared.

